# A demographic approach to understanding the effects of climate on population growth

**DOI:** 10.1101/840512

**Authors:** Nicholas M. Caruso, Christina L. Staudhammer, Leslie J. Rissler

**Author notes:** Author contributions: NMC, CLS, and LJR conceived the idea, NMC developed the models and analyzed the data, and NMC, CLS, and LJR wrote the manuscript.

## Abstract

Amphibian life history traits are affected by temperature and precipitation. Yet, connecting these relationships to population growth, especially for multiple populations within a species, is lacking and precludes our understanding of how amphibians are distributed. Therefore, we constructed Integral Projection Models (IPM) for five populations along an elevational gradient to determine how climate and season affects population growth of a terrestrial salamander *Plethodon montanus* and the importance of demographic vital rates to population growth under varying climate scenarios. We found that population growth was typically higher at the highest elevation compared to the lower elevations whereas varying inactive season conditions, represented by the late fall, winter and early spring, produced a greater variation in population growth than varying active season conditions (late spring, summer, and early fall). Furthermore, survival and growth was consistently more important, as measured by elasticity, compared to fecundity and large females had the greatest elasticity compared to all other sizes. Our results suggest that changing inactive season conditions, especially those that would affect the survival of large individuals, may have the greatest impact on population growth. Therefore, we recommend experimental studies focused on the inactive season to determine the mechanism by which these conditions can affect survival.

---

Climate is a major driver of amphibian life history (Morrison and Hero, 2003) as temperature and precipitation play a key role in amphibian physiological processes (Wells, 2007). Cooler climates often lead to larger body size, larger clutches and eggs, and less frequent clutches among populations of the same species distributed over geographic space (Tilley, 1980; Berven and Gill, 1983; Peterman et al., 2016; reviewed in Morrison and Hero, 2003). Even within populations, warmer temperatures have been associated with reductions in body size (Caruso et al., 2014) or body condition (Reading, 2007), higher survival (Caruso and Rissler, 2019) and lower growth (Muñoz et al., 2016; Caruso and Rissler, 2019). Increased drought frequency can reduce body size (Bendik and Gluesenkamp, 2012) and decrease survival or recruitment (Reading, 2007). While temperature and precipitation can generate broad geographic patterns in life history traits, making connections between the variation in these traits and population growth is a logical next step to understanding how and why species are distributed especially in light of recent and continuing global amphibian declines (Hoffman et al., 2010; Grant et al., 2016).

Understanding how life history traits can contribute to spatial and temporal variation in population growth is a central tenet of life history theory (Stearns, 1977; Caswell, 1982) and important for central questions in ecology, evolution, and conservation science. A mechanistic understanding of how a species’ presence or abundance changes across spatial or temporal gradients can be achieved through understanding the traits that drive population growth (Urban et al., 2016). Likewise, conservation and management of a species requires knowledge of how populations might respond to certain conditions, while developing strategies to maintain or increase abundance are most effective when concentrated on those life stages that have a disproportionately large effect on population growth (Caswell, 2000). Answering these questions requires a demographic modeling approach that can account for how life history traits respond to environmental conditions as well as the sensitivity of population growth to changes in a given life history trait (Caswell, 2001).

Demographic population models have been a useful tool to understand the role of variation in life history (e.g., differences in clutch sizes, age at sexual maturity) on population growth (Caswell, 2001). In a class of models, known as matrix population models, matrices are constructed using stage-specific estimates of vital rates (i.e., survival, fertility, and growth or transitions) that are typically derived from field observations (e.g., life tables, capture-recapture surveys; Caswell, 2001). The eigenvalues and eigenvectors of these projection matrices can be used to estimate population growth as the finite rate of population increase (λ). They can also quantify how proportional changes in a given vital rate can affect population growth through the calculation of vital rate elasticity, a measure that is comparable across populations or species (Caswell, 2000, 2001). Vital rate elasticity has been shown to be a useful tool to compare populations or species: those with more evenly distributed vital rate elasticities are often less likely to decline under environmental stochasticity and typically are of lower conservation concern (Van Allen et al., 2012). These demographic metrics have been used with a variety of organisms such as trees (e.g., Ribeiro do Valle et al., 2007), birds (e.g., Saether and Bakke, 2000), mammals (e.g., Heppell et al., 2000), and amphibians (e.g., Homyack and Haas, 2009), but typically focus on a single population and do not account for intraspecific variation across geographic space (Coulson et al., 2005).

Matrix models are best for species with discrete stages (e.g., egg, juvenile, adult), but may fall short for species that are characterized by continuous state variables (e.g., body size; Easterling et al., 2000). Integral projection models (IPM) are an extension of matrix population models that can provide the same estimates (e.g., population growth, vital rate elasticity) but accommodate continuous state variables (Easterling et al., 2000). Instead of describing survival, growth or transition, and reproduction of stage classes, IPMs use the relationships between these vital rates and a continuous state variable (Easterling et al., 2000) and can be extended to include additional parameters (e.g., climate) that may affect the outcome of vital rates (Ellner and Rees, 2006).

We used IPMs to understand how climate and body size affect population growth of a terrestrial salamander *Plethodon montanus* through their effects on survival, growth, and reproduction. This species is a fully terrestrial montane salamander that is generally found above 1,000 m in elevation within the southern Appalachian Mountains. Like other amphibians, body size, measured as the snout-to-posterior-vent (SVL) has been shown to be an important predictor of survival, growth, and reproduction in this species (Caruso and Rissler, 2019; Caruso and Rissler, *in press*). Moreover, temperature and precipitation are important predictors of plethodontid demography (Caruso and Rissler, 2019) and range limits (Caruso et al., 2019), since these salamanders require cool and moist conditions to exchange gasses across their skin. We combined mark-recapture (Caruso and Rissler, 2019) and museum surveys (Caruso and Rissler, *in press*) to determine the relationships between population vital rates, body size, and climate to parameterize our models. We used IPMs to ask the following questions: 1) How does variation in life history and climate affect population growth, 2) Which vital rates are most important to population growth? and 3) Does vital rate importance vary with climate and season?

## Methods

### Study sites

Our five focal sites were located along an elevational gradient (996 to 1,464 m) in Pisgah National Forest, North Carolina. We chose these sites based on availability of historic collections and the similarity in leaf litter depth, canopy coverage and aspect (Caruso and Rissler, 2019), all of which have been shown to affect terrestrial salamander populations (Pough et al., 1987; Knapp et al., 2004; Wilkins and Peterson, 2000). A reciprocal transplant experiment suggests that the observed differences in demography (Caruso and Rissler, 2019) along this elevational gradient are due predominately to climate (Caruso et al. 2019).

### Demographic data

We combined data from museum (Caruso and Rissler, *in press*) and field surveys (Caruso and Rissler 2019) to obtain estimates of demographic vital rates, and their uncertainty, at all five focal sites. The field data were collected via mark-recapture surveys from 2013 to 2016. During each survey, each unmarked individual was given a unique mark and we measured the length from snout to the posterior margin of the vent (SVL) for all individuals. We used hierarchical Bayesian models to estimate survival and growth using longitudinal capture histories and body size measurements. Survival was estimated using a spatial Cormack-Jolly-Seber model (Schaub and Royle 2014), which accounted for the variation in capture probability and dispersal to estimate survival. Growth rates were estimated using the von Bertalanffy growth curve, which was parameterized for unknown ages (Fabens 1965). Both survival and growth were estimated for two seasons each year (an active season 27 May-13 Oct and an inactive season 14 Oct-26 May), and included parameters for body size, elevation, season, and active season temperature and precipitation. In addition to these parameters, the survival model included parameters for inactive season temperature and snow water equivalent (SWE). The growth model estimated a parameter for standard deviation in growth, and included parameters for asymptotic size for each of the five sites along the elevational gradient. Further details about surveys and models can be found in Caruso and Rissler (2019).

The museum surveys included specimens housed at the US National Museum of Natural History and were collected from 1968 to 1999. For each individual, SVL was measured; reproductive condition was confirmed; and the number of eggs for gravid females was counted via dissection. We used estimates of probability of reproducing using the results from a Bayesian logistic regression, which included parameters for body size and temperature seasonality of the location where each individual was collected. Additionally, these museum data were used to obtain the mean number of eggs per female as an estimate of clutch size. Lastly, we used the results of the hierarchical Bayesian growth model from the skeletochronology analyses to calculate mean recruit size for each focal elevation. This model used a von Bertalanffy (1938) growth curve that was parameterized for known ages. The age of each individual was estimated by the number of lines of arrested growth (LAG) present within the periosteal layer of their long bones, which are formed during annual periods of inactivity (reviewed in Castanet and Smirina, 1990), i.e., in the winter for montane Appalachian salamanders (e.g., Caruso and Rissler, 2019). Lastly, model-based estimates of mean recruit size for each site were determined by using the site-specific growth rates and asymptotic sizes and setting the age to zero. Additional details about methods and models can be found in Caruso and Rissler (*in press*).

### Climate data

We used the DAYMET database (http://www.daymet.org; Thornton et al. 1997) to obtain climate data, utilizing mean maximum temperature (°C) to describe both active and inactive season temperatures. Active and inactive season precipitation variables were characterized by mean precipitation (mm) and snow water equivalent (SWE; kg/m^2^) during each seasonal period, respectively. We used data from 1980 to 2017 to obtain a representation of the range of climate experienced at each site, determining the median and 95% quantiles of each of the four climate variables to parameterize the IPMs (Climate values used can be found in the electronic ESM Figure 1). Because temperature seasonality was found to be associated with the probability of reproduction (Caruso and Rissler, *in press*), we defined this phenomenon following Hijmans et al. (2005), using the WORLDCLIM database and estimating the standard deviation of the mean monthly temperatures at each of our five focal sites.

**Figure 1.**
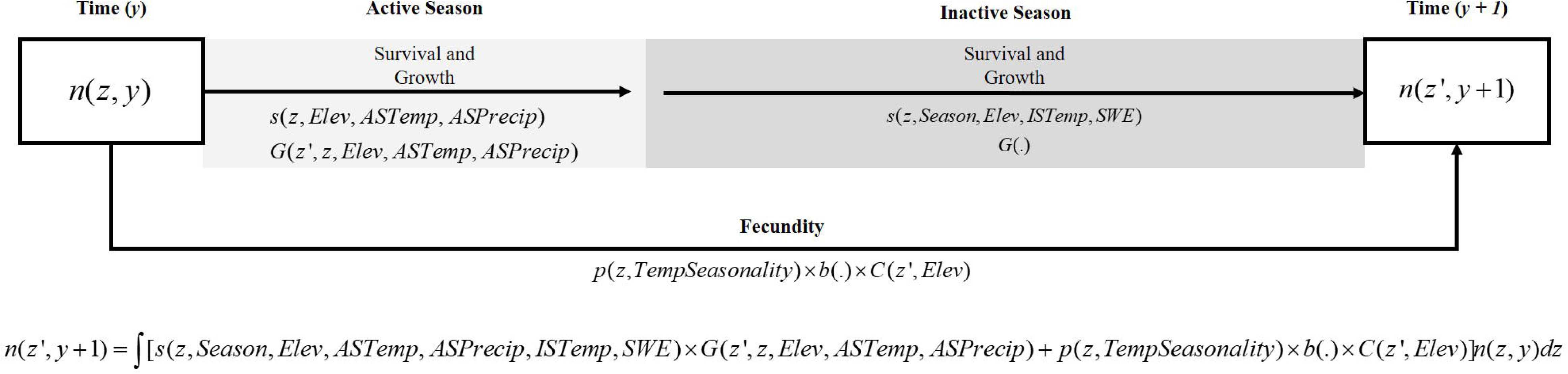
Life cycle diagram and IPM equation. Variables include *n*(*z, y*) = the probability density of *z* at time *y, z* = body size at time *y, z*’ = body size at time *y + 1, s* = survival probability function, *G* = growth kernel, *p* = probability of reproducing, *b* = clutch size, *C* = offspring size, *Elev* = elevation, *ASTemp* = temperature during the active season, *ASPrecip*= precipitation during the active season, *Season* = effect of inactive season compared to active season, *ISTemp* = Inactive season temperature, *SWE* = inactive season snow water equivalent, and *TempSeasonality* = temperature seasonality.

### IPM

We used the estimates of demographic vital rates, their uncertainty, and climate data to parameterize site-specific and climate-varying deterministic post-reproductive census IPMs. As IPMs estimate population dynamics at discrete time points using individual-level continuous state variables, we defined our IPMs using body size (as measured by SVL in mm), and our discrete time step was defined as a year. Therefore, for each site our IPM describes *z’*, the body size distribution at time *y +* 1, given *z*, the body size distribution at time *y*, as a function of survival, growth and fecundity (Fig. 1). The function *n* describes the population size distribution function, *s* is probability of survival, *G* is the growth function, *p* is the probability of reproducing, *b* is clutch size, and *C* is the recruitment size distribution.

Survival probability was estimated as a function of body size, elevation of each site (*Elev*), and the elevation-specific active season temperature (*ASTemp*_*Elev*_), active season precipitation (*ASPrecip*_*Elev*_), inactive season temperature (*ISTemp*_*Elev*_) and inactive season SWE (*SWE*_*Elev*_; Table 1). The growth function was estimated using the Von Bertalanffy growth equation as a function of the body size distribution at time *y +* 1 given the body size at time *y*, elevation, and the elevation-specific active season temperature and active season precipitation. The size probability density function of individuals in time *y +* 1 was described as a normal probability density function, where the mean was the expected size in time *y +* 1 and the standard deviation *σ*_*G*_ that was obtained from the results of the growth model (Table 1). As both survival and growth were estimated for active and inactive seasons separately, we weighted each by the number of days for each season (138 and 227 respectively; Table 1).

**Table 1.**
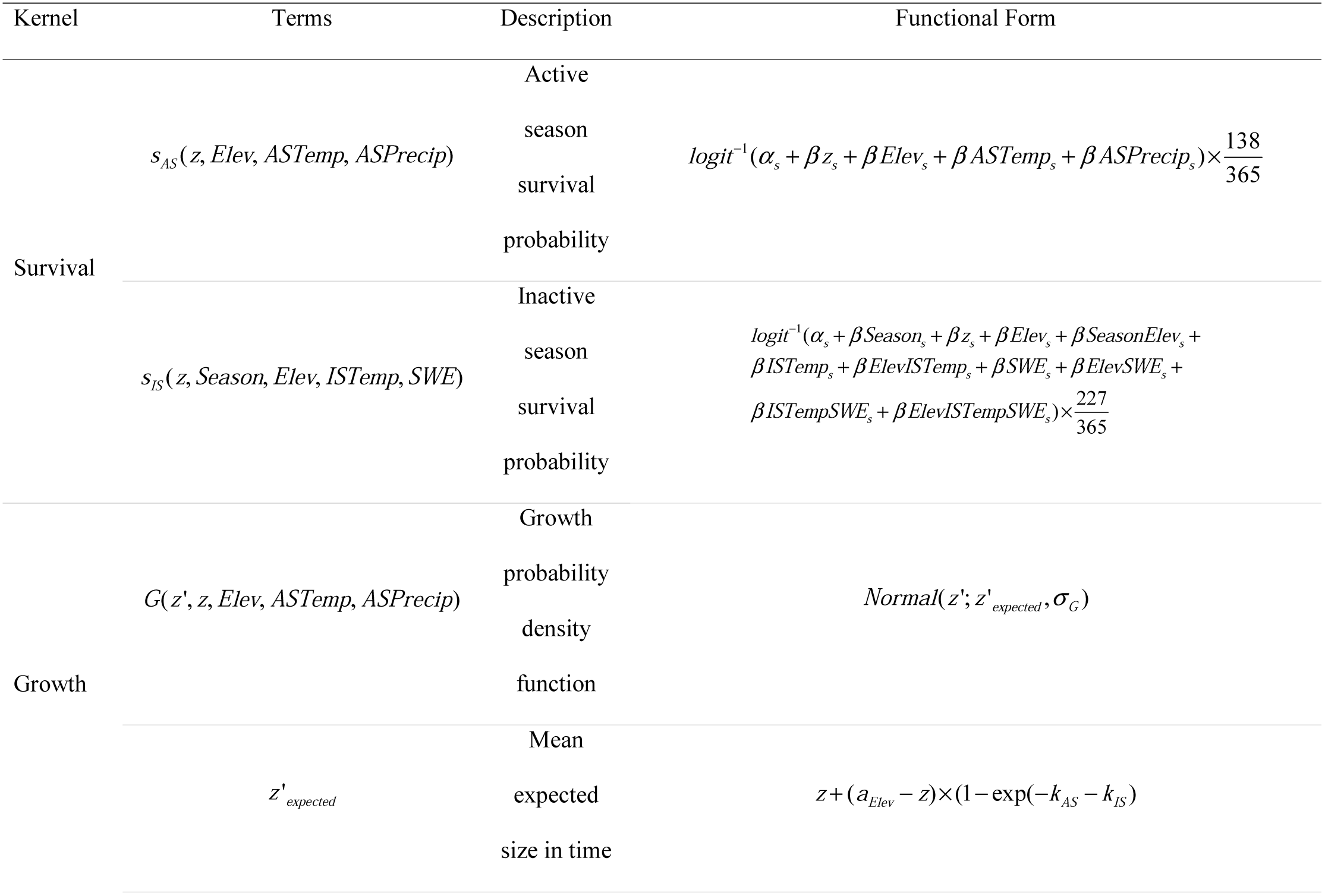
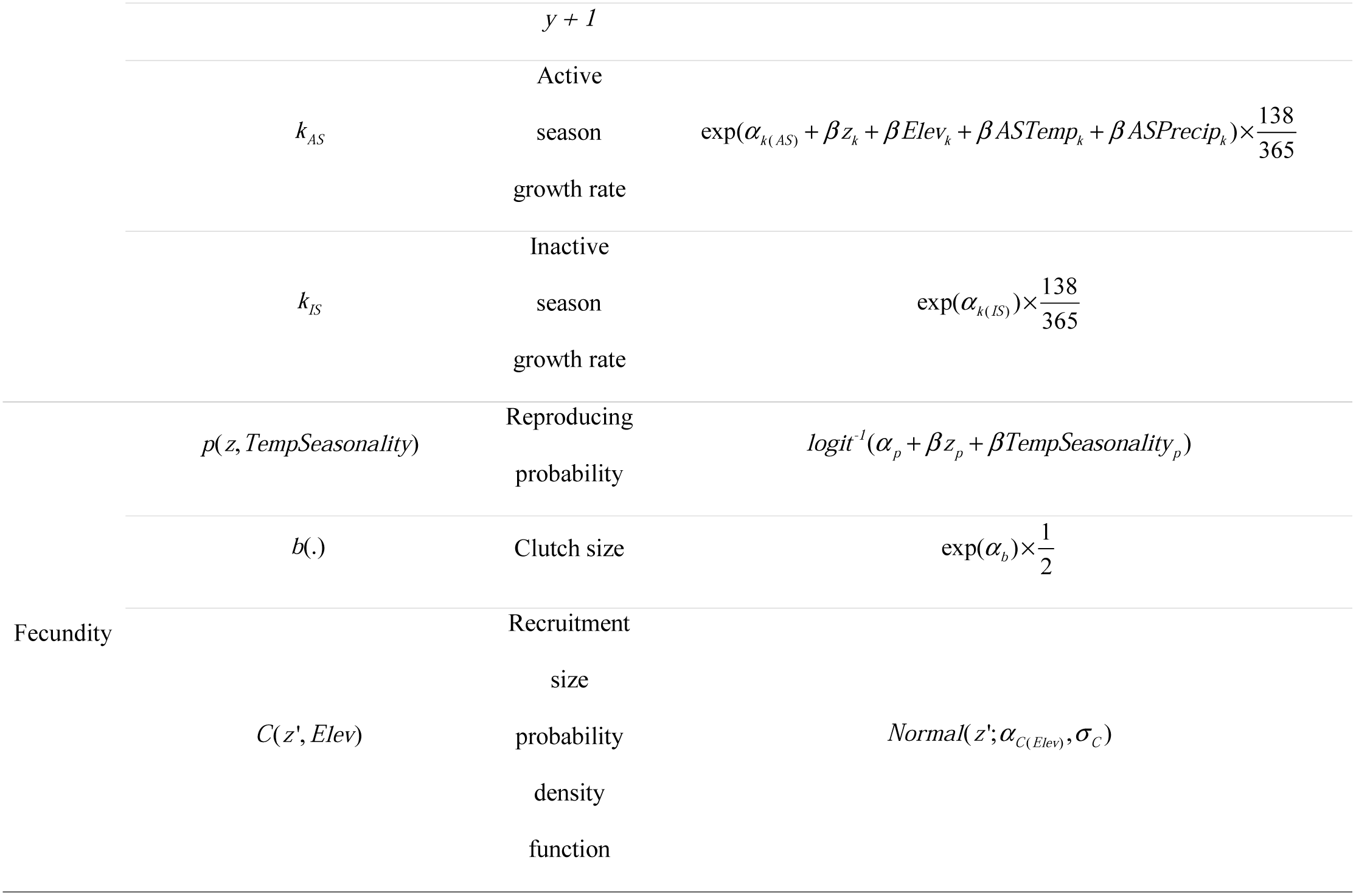
Components and parameter terms for the IPM model. Variables include *z* = body size in time *y, z*’ = body size at time *y* + 1, *Elev* = elevation, *ASTemp* = temperature during the active season, *ASPrecip* = precipitation during the active season, *Season* = effect of inactive season compared to active season, *ISTemp* = Inactive season temperature, *SWE* = inactive season snow water equivalent, and *TempSeasonality* = temperature seasonality. In the functional form, *α* denotes intercept, *β* is a fixed effect for a given variable, *a*_*Elev*_ is the elevation specific asymptotic size, *σ* = is the standard deviation.

The probability of reproducing was estimated as a function of body size (*z*) and temperature seasonality (*TempSeasonality*) of each site. Clutch size was estimated as the mean number eggs per female multiplied by 0.5, which assumes a 1:1 sex ratio of the offspring (Caruso and Rissler, *in press*; Table 1). Lastly, we describe the elevation-specific recruitment size probability density function during time *y +* 1 using a normal probability density function where the mean is elevation-specific mean offspring size and a standard deviation of one (Table 1).

### Simulating population dynamics with climate

For each of our five focal sites, we fit IPMs exploring the effect of both active season and inactive season conditions. We explored the effects of each season by fitting IPMs with all combinations of high, mid, and low temperature and precipitation (or SWE) values within a season while conditions in the other season were held at their median values (i.e., nine climate combinations per season for each of the five sites). For example, to investigate how active season conditions affect population growth, one of the nine climate combinations included low active season temperature and high active season precipitation, and both inactive season temperature and SWE were held at their median values. For each of the five focal sites we had 17 climate combinations (85 total site-climate combinations), since climate combinations with all median values were duplicated while exploring both active and inactive season conditions (i.e., median active season temperature and precipitation, median inactive season temperature and SWE).

For each site-climate combination, we explored the effect of parameter uncertainty by randomly drawing a value for each parameter from their respective posterior distributions (see Table 1) and replicating this process independently 1,000 times. We used each IPM simulation to calculate the finite rate of population increase (*λ*), and mean elasticity surface of *λ* to changes in the kernel for all simulations (Easterling et al., 2000). For each site and climate combination, we calculated the mean elasticity surface of all simulations as well as the elasticity of *λ* to both survival-growth and fecundity kernels (Easterling et al., 2000). Lastly, we used the mean elasticity surfaces to calculate elasticity evenness, in which high evenness values (i.e., closer to one) indicate a population or species with more symmetrical importance across demographic rates while evenness values closer to zero indicate more asymmetric importance (Van Allen et al., 2012). We used Program R (version 3.4.4; R Core Team, 2018) for all analyses and all code and data can be found at https://github.com/PlethodoNick/Pmontanus-IPM.

## Results

Estimates of *λ* were generally greater at the highest elevation compared to the lower elevations at most temperature and precipitation conditions tested (Figs. 2, 3). Within a given elevation, the distributions of *λ* showed little variation across the range of active season temperature and precipitation combinations (Fig. 2), whereas altering inactive season conditions resulted in a greater range of *λ* values (Fig. 3; ESM Table 1). Generally, *λ* increased with increasing active season precipitation, whereas increasing temperature did not always lead to an increase in *λ*, especially at lower elevations (Fig. 2; ESM Table 1). Inactive season conditions were relatively more complex and were elevation-specific. At the two lowest elevations, *λ* increased with increasing temperature but did not show a trend with SWE (Fig. 3; ESM Table 1). Likewise at the mid elevation, inactive season temperature and *λ* showed a stronger relationship than SWE and *λ*, however at this elevation lower temperatures led to higher *λ* (Fig. 3; ESM Table 1). At the next highest elevation (1,300), increasing SWE led to higher *λ* but at the highest elevation site-specific values of *λ* did not show patterns with either inactive season temperature or SWE (Fig. 3; ESM Table 1).

**Figure 2.**
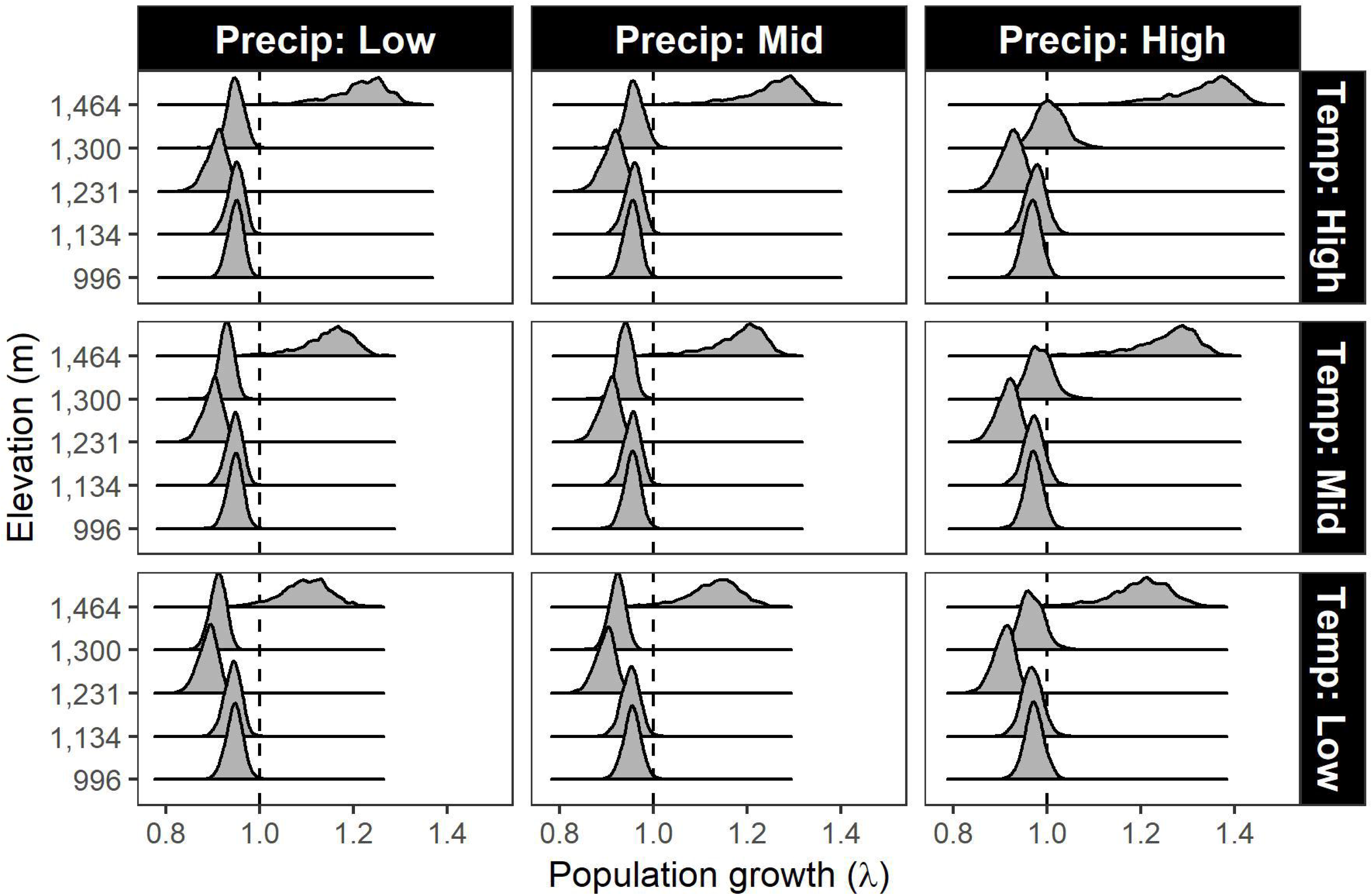
Distribution of *λ* for each focal elevation and with the nine climate scenarios for changing active season conditions. Dashed line shows *λ* of one, indicating a population that remains the same.

**Figure 3.**
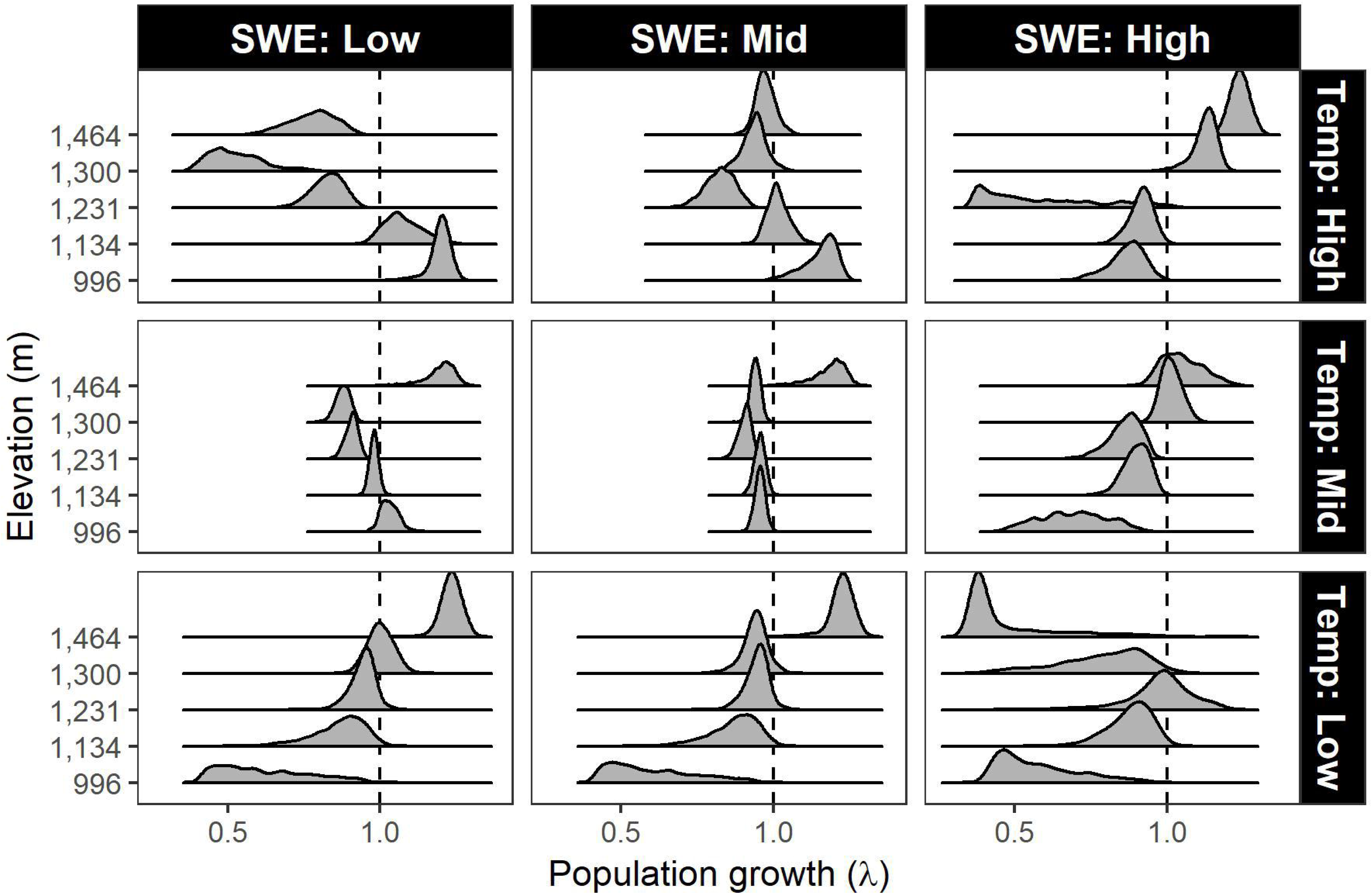
Distribution of *λ* for each focal elevation and with the nine climate scenarios for changing inactive season conditions. Dashed line shows *λ* of one, indicating a population that remains the same.

Elasticity analyses show that the survival-growth kernel (mean elasticity = 0.879 – 0.999) consistently had a greater importance than fecundity kernel (mean elasticity = 0.001 – 0.121) to population growth. For all sites and climate conditions, we found that survival of large females was most important to population growth with elasticity values generally decreasing towards the smaller size classes (Figs. 4, 5), although the highest elevation site typically had more evenly distributed elasticities compared to the lower elevations (Figs. 4, 5, 6). Our elasticity results also demonstrate the relatively low importance of fecundity (represented in the bottom right side of each panel in figs. 4 and 5) compared to survival and growth. Lastly, we found that for all sites average *λ* generally increased with increasing elasticity evenness (Fig. 6). Therefore, climate conditions that typically led to higher population growth also led to an increase in the importance of fecundity to population growth and more even elasticities across all body sizes.

**Figure 4.**
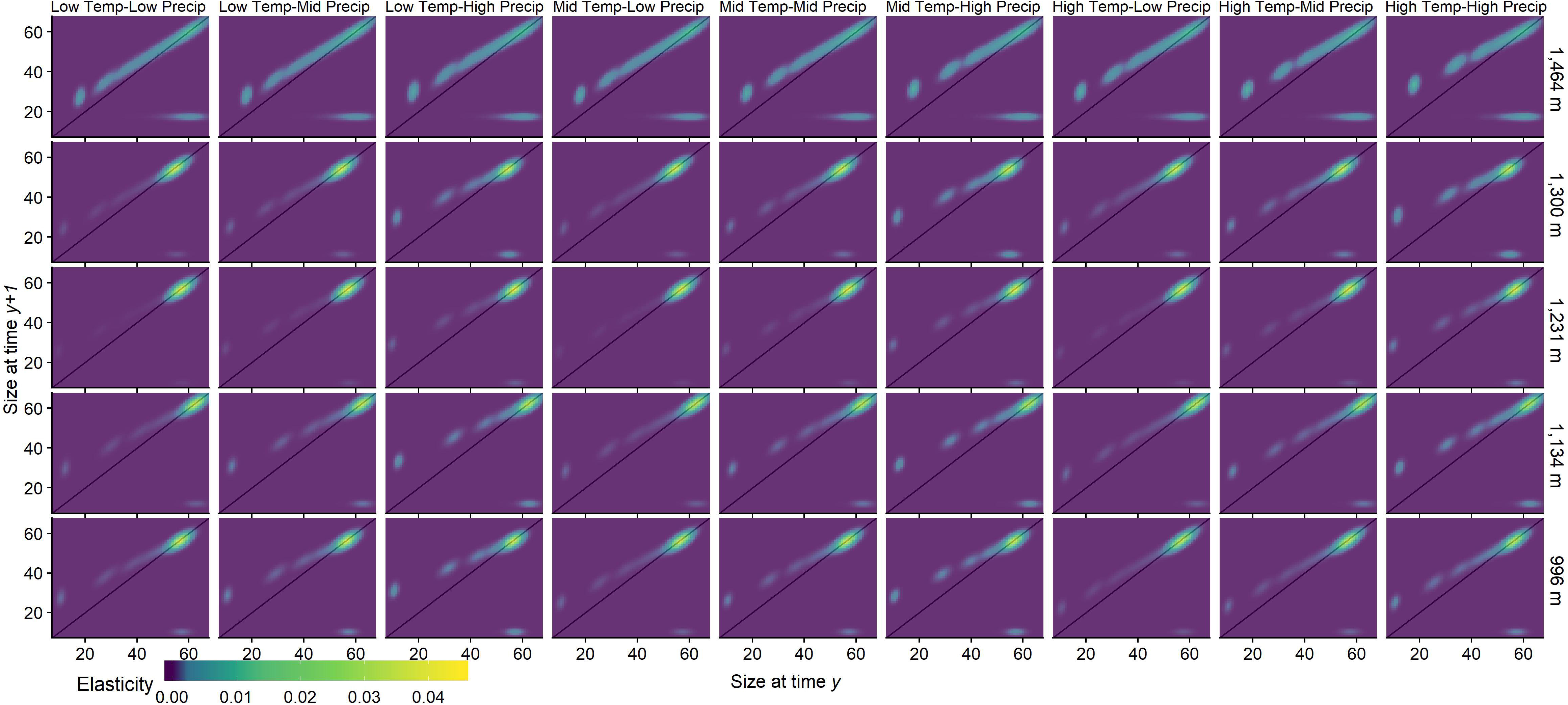
Elasticity of vital rates for each elevation and the nine climate scenarios for changing active season conditions. Each pixel represents the elasticity value for an individual of size *z’* at time *y +* 1 (y-axis) given their size (*z*) at time *y* (x-axis). The solid line indicates individuals that survive but remain the same size from time *y* to time *y +* 1. Darker colors denote lower elasticity while lighter colors show high elasticity.

**Figure 5.**
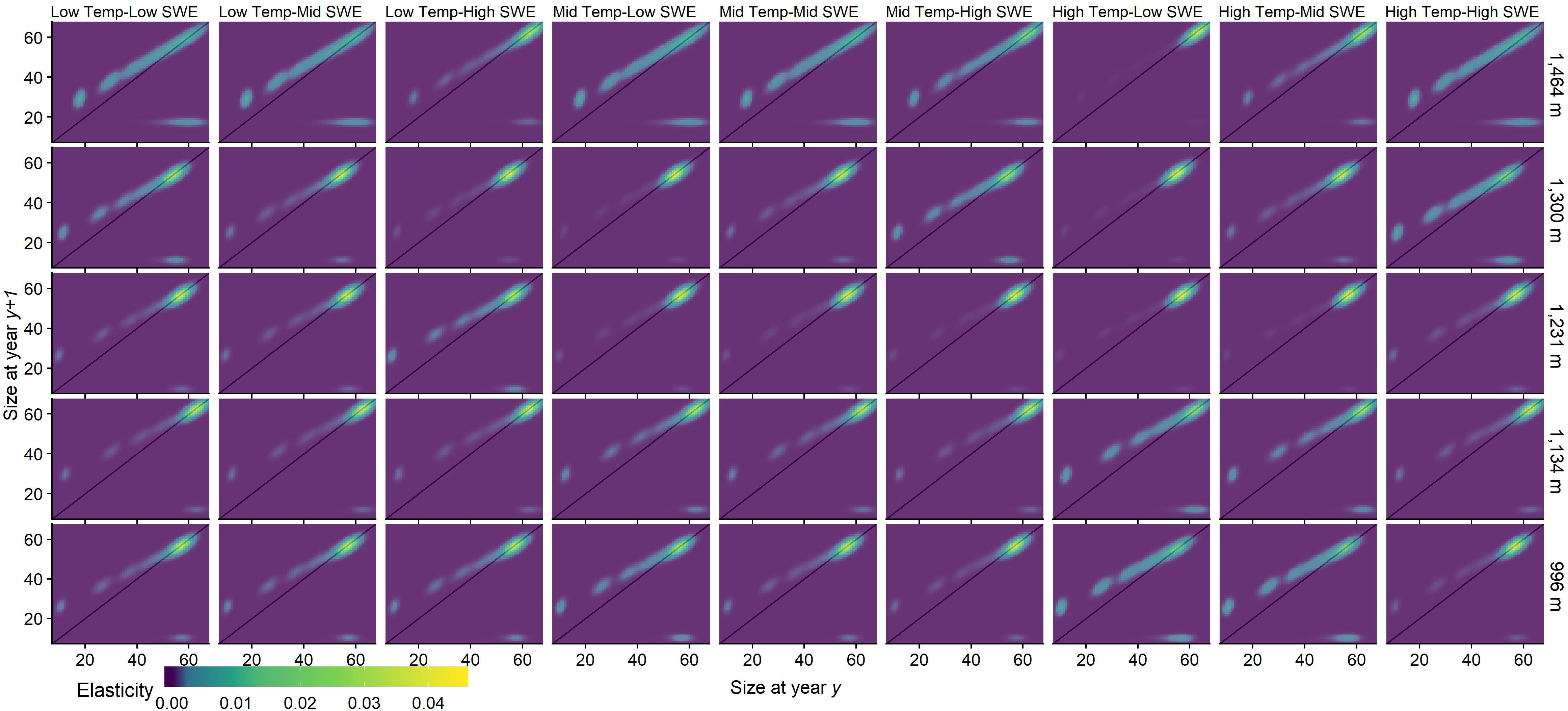
Elasticity of vital rates for each elevation and the nine climate scenarios for changing inactive season conditions. Each pixel represents the elasticity value for an individual of size *z’* at time *y +* 1 (y-axis) given their size (*z*) at time *y* (x-axis). The solid line indicates individuals that survive but remain the same size from time *y* to time *y +* 1. Darker colors denote lower elasticity while lighter colors show high elasticity.

**Figure 6.**
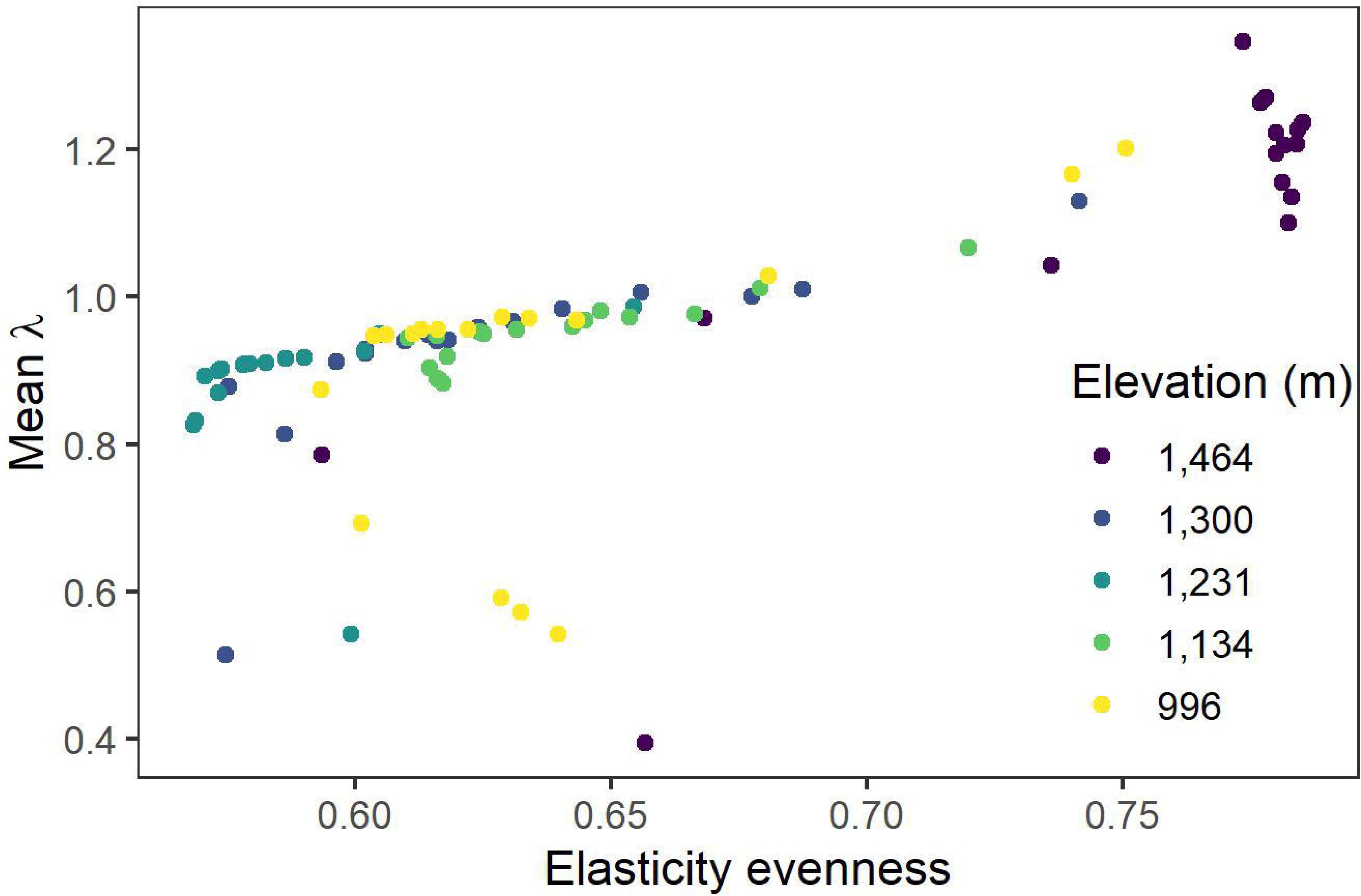
The relationship between median *λ* and elasticity evenness. Each point represents a single elevation and climate scenario combination. Colors denote site elevation with darker colors as higher elevations.

## Discussion

We used an IPM approach to demonstrate how climate can affect population growth in *Plethodon montanus* across an elevational gradient by combing field and museum estimates of survival, growth, and reproduction. Our results demonstrate that inactive season conditions had a disproportionately large effect on population growth compared to the conditions in the active season. Life stages were generally uneven in their importance to population growth and the survival of large females was the most important vital rate for *P. montanus* across all elevations. However, at the highest elevation, the evenness of vital rate elasticity to population growth was higher compared to other elevations. The utility of these models extends beyond understanding the current distribution of *P. montanus* by providing insight into how changes in climate can affect these populations.

### Lower elevation range limits

Recent evidence suggests that the warmer edge (i.e., lower elevation or latitude) of amphibian ranges are limited by climate (Gifford and Kozak 2012; Cunningham et al. 2016; Lyons et al. 2016; Grant et al., 2018; Caruso et al., 2019). While we did not formally test this hypothesis, our results support these studies. Along our elevational gradient, population growth, driven by climate, was typically lower at the low elevations compared to the higher elevation. It is important to note that while some of our median estimates of *λ* for different climate scenarios were less than one, which indicates a population that is below replacement, our results do not suggest that populations are declining. Our prospective perturbation analysis provides a “what if” scenario for how changes in vital rates could affect population growth; a retrospective perturbation analysis would be required to determine actual population trajectories (Caswell 2000). Furthermore, providing a more realistic scenario for these populations would require estimates of the variance, covariance, and density-dependence of vital rates (Benton et al. 1995; Gaillard et al. 1998). For example, a low-density population would have higher rates of survival compared to a higher-density population due to intraspecific competition. Lastly, models that exclude dispersal can underestimate population growth (Tavecchia et al. 2016); however, dispersal has been shown to be limited in terrestrial plethodontids (Liebgold et al. 2011; Caruso and Rissler, 2019).

### Vital rate importance

Higher importance of survival, especially of the largest or oldest individuals or stage is typical of long-lived species (Heppell et al. 2000), and our results are consistent with this pattern. Although few studies deploy demographic models (e.g., matrix models) for plethodontid species, high importance of the largest size or class of females is typical (e.g., Homyack and Haas, 2009; Lindström et al., 2010), and is for other salamander species as well (Schmidt et al., 2005). These results suggest that changes in survival of these size or stage classes will have a disproportionately large effect on population growth compared to other vital rates, and it is the most important vital rate for population persistence. Because of high elasticity, life history theory predicts that selection will act upon traits that are associated with survival of large adults (Benton et al. 1995; Benton and Grant 1999; Saether and Bakke 2000), such as physiological acclimatization and behavioral avoidance to adverse environmental conditions (Riddell et al., 2018). Elasticity analyses suggest that management techniques that increase survival of large females (e.g., availability of surface retreats; Rittenhouse et al., 2008; Otto et al., 2014) will have the greatest effect on population growth.

Elasticity evenness is a useful metric of the distribution of vital rate importance and has been shown to be associated with species’ conservation status and the susceptibility of populations to decline (Van Allen et al., 2012). We found that population growth generally increased with increasing elasticity evenness at all sites, but the highest elevation site typically had the most evenly distributed elasticity compared to the rest of the elevational gradient (Fig. 5). Because populations with low elasticity evenness are more likely to be negatively affected by environmental perturbations that reduce survival or fecundity (Van Allen et al., 2012), we would expect populations at higher elevations to have less variability in population growth compared to those at lower elevations.

### Importance of inactive season

Inactive season conditions have been shown to be important to population growth in amphibians (Benard, 2015; Muths et al., 2017). We found that the inactive season produced a greater variation in population growth compared to the active season (Figs. 2, 3) at all elevations. This could result from a larger range of climate variables included in our models during the inactive season compared to the active season (ESM Fig 1). However, even when we limit the inactive season conditions to their low and average values, the response of population growth to those varying conditions is still greater than the response to the full range of active season conditions (Figs., 2, 3, ESM Table 1).

The importance of inactive season survival may mean even greater future negative consequence to these populations because winter temperatures in the study region continue to increase at a greater rate compared to summer temperatures (Xia et al., 2014). Although our results show that warmer inactive season temperatures or reductions in SWE do not always lead to reductions in population growth (Fig. 3; ESM Table 1), warmer temperatures that reduce snowpack would subject soils, and potentially salamanders, to freezing temperatures (Decker et al., 2003; Henry 2008; Bale and Hayward 2010; Brown and DeGaetano 2011).

Climate change is predicted to substantially decrease environmental suitability for Southern Appalachian plethodontids (Milanovich et al. 2010; Sutton et al. 2015), but these predictions may bias losses by not accounting for physiological adaptations (Riddell et al., 2018) or a mechanistic understanding of how climate affects demography and population growth (Buckley et al., 2010; Urban et al., 2016). Here we show that changes in the inactive season climate can generate a greater variation in population growth compared to variation during the active season. Moreover, temperature and precipitation were not equally influential to population growth within a season. During the active season, population growth was influenced primarily by precipitation, whereas temperature was more important during the inactive season. Our results show that survival of large females is the most important vital rate for *P. montanus*. Therefore, future studies should identify the mechanisms (e.g., reduced snowpack) that can increase or decrease survival of this stage to better predict how populations with respond to changes in climate.

## Supporting information

Electronic Supplemental Material

## Acknowledgements

Thanks to C. Houser, and E. Kabay for assistance in the field, J. Jacobs, R. McDiarmid, A. Wynn, K. Tighe, and S. Gotte for assistance with museum specimens and to H. Wimer and K. Lackey for assistance with skeletochronology.

## Ethical approval

All applicable institutional and/or national guidelines for the care and use of animals were followed.

