## Supplementary material for "A demographic approach to understanding the effects of climate on population growth": Electronic Supplemental Material

**ESM Figure 1.** Climate values used in the IPMs for each site. Triangles represent the median, while the two squares are the upper and lower 95% quantiles. Data were obtained from the DAYMET database (http://www.daymet.org; Thornton et al. 1997)

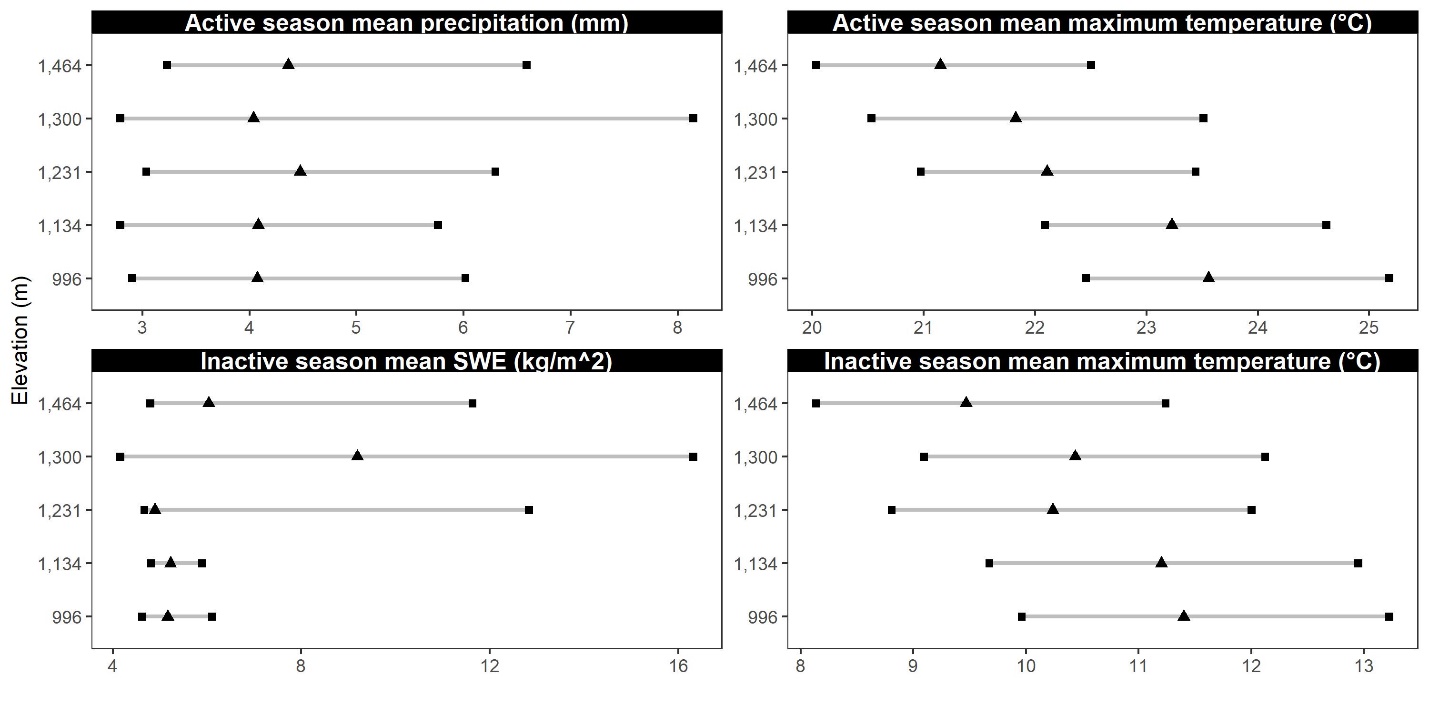

**ESM Table 1.** Mean, standard deviation, and 95% quantiles for population growth (λ) for each site and climate combination.

| Site  (Elevation) | Season | Temperature | Precipitation | Mean  λ | Standard  Deviation | Upper  99% CI | Lower  99% CI |
| --- | --- | --- | --- | --- | --- | --- | --- |
| SPG  (996) | Active | High | Low | 0.95 | 0.02 | 0.98 | 0.92 |
|  |  |  | Mid | 0.95 | 0.02 | 0.99 | 0.92 |
|  |  |  | High | 0.97 | 0.02 | 1.00 | 0.93 |
|  |  | Mid | Low | 0.95 | 0.02 | 0.98 | 0.91 |
|  |  |  | Mid | 0.95 | 0.02 | 0.99 | 0.92 |
|  |  |  | High | 0.97 | 0.02 | 1.01 | 0.93 |
|  |  | Low | Low | 0.95 | 0.02 | 0.98 | 0.91 |
|  |  |  | Mid | 0.95 | 0.02 | 0.99 | 0.92 |
|  |  |  | High | 0.97 | 0.02 | 1.01 | 0.93 |
|  | Inactive | High | Low | 1.20 | 0.04 | 1.25 | 1.08 |
|  |  |  | Mid | 1.15 | 0.06 | 1.23 | 1.02 |
|  |  |  | High | 0.86 | 0.06 | 0.97 | 0.72 |
|  |  | Mid | Low | 1.03 | 0.03 | 1.09 | 0.98 |
|  |  |  | Mid | 0.95 | 0.02 | 0.99 | 0.92 |
|  |  |  | High | 0.69 | 0.11 | 0.89 | 0.49 |
|  |  | Low | Low | 0.62 | 0.15 | 0.92 | 0.42 |
|  |  |  | Mid | 0.60 | 0.14 | 0.91 | 0.42 |
|  |  |  | High | 0.57 | 0.13 | 0.87 | 0.42 |
| IMG  (1,134) | Active | High | Low | 0.95 | 0.02 | 0.98 | 0.91 |
|  |  |  | Mid | 0.96 | 0.02 | 0.99 | 0.92 |
|  |  |  | High | 0.98 | 0.02 | 1.02 | 0.93 |
|  |  | Mid | Low | 0.95 | 0.02 | 0.98 | 0.91 |
|  |  |  | Mid | 0.95 | 0.02 | 0.99 | 0.92 |
|  |  |  | High | 0.97 | 0.02 | 1.01 | 0.93 |
|  |  | Low | Low | 0.94 | 0.02 | 0.97 | 0.91 |
|  |  |  | Mid | 0.95 | 0.02 | 0.99 | 0.91 |
|  |  |  | High | 0.97 | 0.02 | 1.01 | 0.93 |
|  | Inactive | High | Low | 1.08 | 0.06 | 1.21 | 0.97 |
|  |  |  | Mid | 1.01 | 0.04 | 1.09 | 0.95 |
|  |  |  | High | 0.91 | 0.04 | 0.98 | 0.82 |
|  |  | Mid | Low | 0.98 | 0.01 | 1.01 | 0.95 |
|  |  |  | Mid | 0.95 | 0.02 | 0.99 | 0.92 |
|  |  |  | High | 0.90 | 0.04 | 0.97 | 0.80 |
|  |  | Low | Low | 0.87 | 0.09 | 1.00 | 0.66 |
|  |  |  | Mid | 0.87 | 0.08 | 0.99 | 0.68 |
|  |  |  | High | 0.88 | 0.07 | 0.99 | 0.71 |
| HG  (1,231) | Active | High | Low | 0.91 | 0.02 | 0.94 | 0.86 |
|  |  |  | Mid | 0.91 | 0.02 | 0.95 | 0.86 |
|  |  |  | High | 0.92 | 0.03 | 0.97 | 0.87 |
|  |  | Mid | Low | 0.90 | 0.02 | 0.93 | 0.85 |
|  |  |  | Mid | 0.91 | 0.02 | 0.94 | 0.86 |
|  |  |  | High | 0.92 | 0.02 | 0.96 | 0.87 |
|  |  | Low | Low | 0.89 | 0.02 | 0.92 | 0.85 |
|  |  |  | Mid | 0.90 | 0.02 | 0.93 | 0.85 |
|  |  |  | High | 0.91 | 0.02 | 0.95 | 0.86 |
|  | Inactive | High | Low | 0.83 | 0.05 | 0.92 | 0.71 |
|  |  |  | Mid | 0.82 | 0.05 | 0.91 | 0.71 |
|  |  |  | High | 0.60 | 0.20 | 0.99 | 0.37 |
|  |  | Mid | Low | 0.91 | 0.02 | 0.94 | 0.86 |
|  |  |  | Mid | 0.91 | 0.02 | 0.94 | 0.86 |
|  |  |  | High | 0.86 | 0.05 | 0.94 | 0.74 |
|  |  | Low | Low | 0.94 | 0.04 | 1.01 | 0.85 |
|  |  |  | Mid | 0.94 | 0.04 | 1.01 | 0.86 |
|  |  |  | High | 0.98 | 0.11 | 1.16 | 0.70 |
| BBT  (1,300) | Active | High | Low | 0.95 | 0.02 | 0.98 | 0.91 |
|  |  |  | Mid | 0.96 | 0.02 | 1.00 | 0.92 |
|  |  |  | High | 1.01 | 0.03 | 1.08 | 0.95 |
|  |  | Mid | Low | 0.93 | 0.02 | 0.96 | 0.90 |
|  |  |  | Mid | 0.94 | 0.02 | 0.97 | 0.91 |
|  |  |  | High | 0.98 | 0.03 | 1.04 | 0.93 |
|  |  | Low | Low | 0.91 | 0.02 | 0.94 | 0.88 |
|  |  |  | Mid | 0.92 | 0.02 | 0.95 | 0.89 |
|  |  |  | High | 0.97 | 0.03 | 1.02 | 0.92 |
|  | Inactive | High | Low | 0.53 | 0.09 | 0.75 | 0.39 |
|  |  |  | Mid | 0.94 | 0.04 | 1.02 | 0.85 |
|  |  |  | High | 1.12 | 0.04 | 1.18 | 1.01 |
|  |  | Mid | Low | 0.88 | 0.03 | 0.92 | 0.82 |
|  |  |  | Mid | 0.94 | 0.02 | 0.97 | 0.91 |
|  |  |  | High | 1.01 | 0.03 | 1.08 | 0.96 |
|  |  | Low | Low | 1.00 | 0.04 | 1.08 | 0.93 |
|  |  |  | Mid | 0.94 | 0.04 | 1.02 | 0.85 |
|  |  |  | High | 0.79 | 0.14 | 0.99 | 0.47 |
| CG  (1,464) | Active | High | Low | 1.21 | 0.06 | 1.30 | 1.05 |
|  |  |  | Mid | 1.25 | 0.07 | 1.34 | 1.08 |
|  |  |  | High | 1.33 | 0.08 | 1.43 | 1.14 |
|  |  | Mid | Low | 1.15 | 0.05 | 1.23 | 1.01 |
|  |  |  | Mid | 1.18 | 0.06 | 1.26 | 1.04 |
|  |  |  | High | 1.26 | 0.07 | 1.35 | 1.09 |
|  |  | Low | Low | 1.10 | 0.05 | 1.19 | 0.99 |
|  |  |  | Mid | 1.13 | 0.05 | 1.23 | 1.00 |
|  |  |  | High | 1.20 | 0.07 | 1.31 | 1.04 |
|  | Inactive | High | Low | 0.78 | 0.08 | 0.91 | 0.61 |
|  |  |  | Mid | 0.97 | 0.03 | 1.05 | 0.91 |
|  |  |  | High | 1.24 | 0.03 | 1.29 | 1.18 |
|  |  | Mid | Low | 1.19 | 0.06 | 1.27 | 1.05 |
|  |  |  | Mid | 1.18 | 0.06 | 1.26 | 1.04 |
|  |  |  | High | 1.05 | 0.07 | 1.19 | 0.94 |
|  |  | Low | Low | 1.23 | 0.03 | 1.29 | 1.17 |
|  |  |  | Mid | 1.22 | 0.04 | 1.29 | 1.11 |
|  |  |  | High | 0.47 | 0.15 | 0.90 | 0.35 |
